# Two parallel pathways implement robust propionate catabolism and detoxification in mycobacteria

**DOI:** 10.1101/258947

**Authors:** Katja Tummler, Michael Zimmermann, Olga T. Schubert, Ruedi Aebersold, Clemens Kühn, Uwe Sauer, Edda Klipp

## Abstract

Tuberculosis remains a major global health threat with over 1.5 million deaths each year. Mycobacterium tuberculosis’ success story is related to a flexible metabolism, allowing growth despite restrictive conditions within the human host.

Host lipids stores are a major carbon source in vivo. Their catabolism yields propionyl-CoA, which is processed by two parallel pathways, the methylmalonyl CoA pathway and the methylcitrate pathway. Both pathways are considered potential drug targets. The methylcitrate pathway is upregulated in the pathological context. However, intermediates of this pathway can be cytotoxic and Mtb’s preference for its usage remains unclear.

We combine thermodynamic kinetic modeling, quantitative proteomics and time-resolved metabolomics to characterize the interplay between the two pathways and to show their functionalities in an efficient and fast propionate catabolism.

We find that the methylcitrate pathway acts as a transcriptionally regulated, high capacity catabolic pathway due to its favorable thermodynamics and metabolic control distribution. In contrast, the methylmalonyl pathway is constitutively fulfilling biosynthetic tasks and can quickly detoxify propionate pulses, but is thermodynamically restricted to lower capacity.

## Introduction

Tuberculosis is the bacterial disease with the highest human death toll every year [47]. A highly flexible and adaptive metabolism is thought to be a major pathogenic trait of the causative agent *Mycobacterium tuberculosis* (*Mtb*) [32, 35]. Accordingly pathways required for the catabolism of host-derived carbon sources are promising targets for drug development [5, 27, 45], for which, however, in depth mechanistic understanding is required.

*Mtb* resides within the hostile and ever-changing environment of the host phagosome, which is naturally poor in nutrients. Recent evidence suggests that the intracellular bacteria likely feed on substrates containing two or three carbon atoms [5], the end products of fatty acid and cholesterol catabolism [29, 45, 49].In *Mtb*, two parallel pathways are responsible for the catabolism of such compounds, the methylcitrate (MCIT) and the methylmalonyl CoA (MMCO) pathway (Figure 1). The two seemingly redundant pathways can both fulfill the task of fueling the central carbon metabolism from three-carbon compounds. However, they employ a different set of biochemical conversions and show a distinct expression pattern in experimental studies. The reason why both pathways are retained in the bacteria and the conditions under which either pathway is used preferentially have been unknown and are the subject of this study.

**Figure 1:**
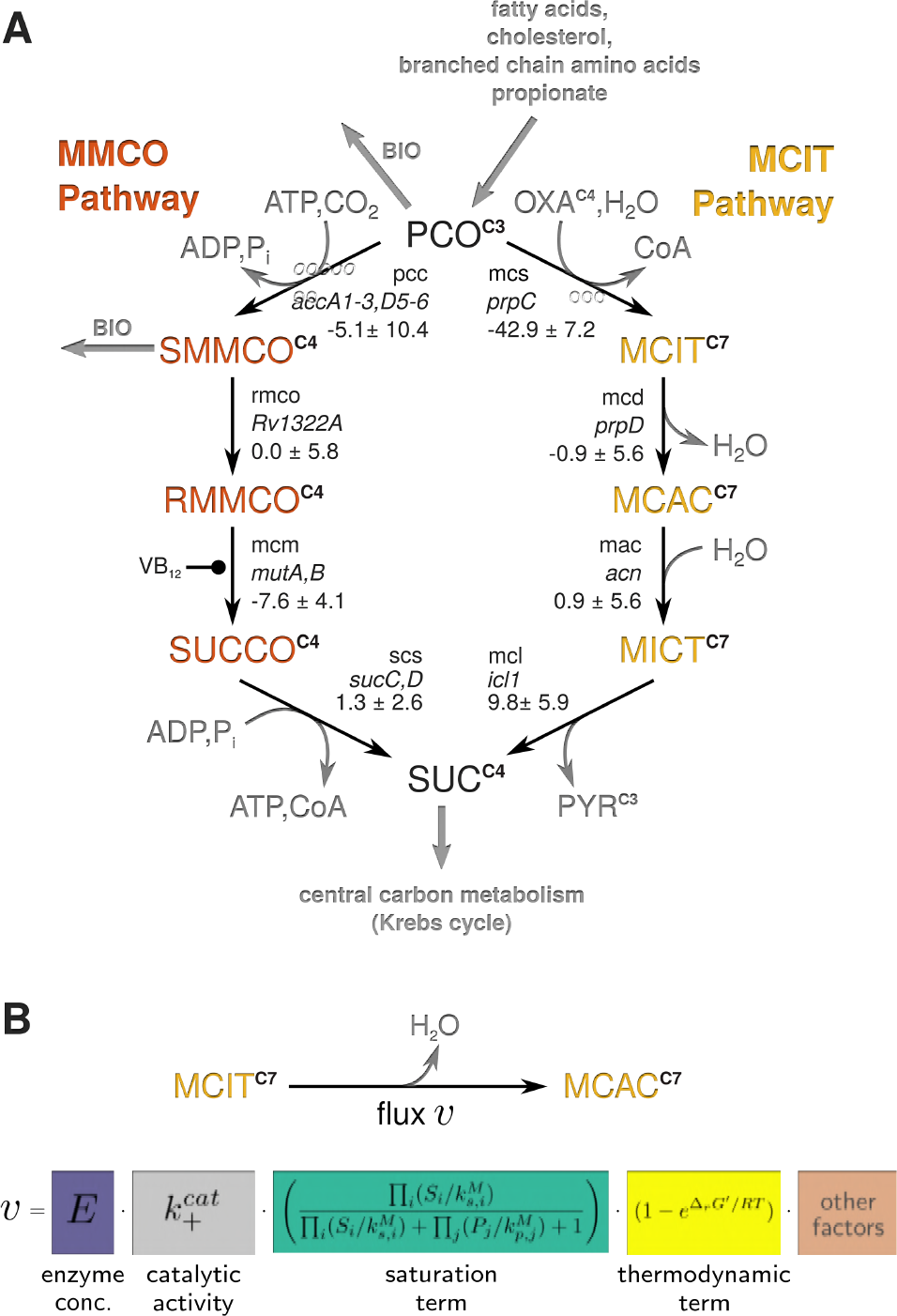
Pathways for propionate catabolism. **A** Schematic representation of the two pathways studied. Each reaction is depicted by an arrow and labeled with the reaction name, the corresponding gene name(s) in italics, and their standard Gibbs energy ∆*_r_G*^0^ in kJ/mol (see also Supplementary Table S2). Gray arrows depict uptake reactions and network derived biomass fluxes. Intermediates of the MCIT pathway are shown in yellow, of the MMCO pathway in orange. Boundary species (cofactors and pathway endpoints) are shown in gray. Superscripts denote the number of carbons for each metabolite, neglecting the carbon in cofactor A. Abbreviations: CoA - cofactor A, PCO - propionyl-CoA, S/RMMCO - S/R-methylmalonyl-CoA, SUCCO - succinyl-CoA, SUC - succinate, OXA - oxaloacetate, MCIT - methylcitrate, MCAC - methylaconiate, MICT - methylisocitrate, VB_12_ - vitamin B_12_ (see also Supplementary Table S1.) **B** Major factors defining the amount of flux through a reaction [25]. Other factors, which are not covered by the models here can be regulatory processes such as post-translational modifications or allosteric regulation.

The MCIT pathway, in analogy to the glyoxylate cycle in the central carbon metabolism, invests one oxaloacetate molecule to convert the three-carbon compound propionyl-CoA (PCO) to pyruvate under the release of succinate, which feed into lower glycolysis and the Krebs cycle, respectively. Enzymes of the pathway are up-regulated and essential in the pathogenic context [4, 13, 15, 21, 34, 37, 38, 41]. Despite its preferential usage in the bacteria, the MCIT pathway also comprises some risks. Its intermediates, especially methylcitrate, are toxic at high concentrations [43, 45]. This toxicity is linked to the inhibitory effect on fructose 1,6-bisphosphatase in the central carbon metabolism [10].

The MMCO pathway depends on ATP activation in the first reaction step, thereby allowing for the additional fixation of one molecule CO_2_. The ATP is regained in the last step of the pathway that again produces succinate. Furthermore, the pathway fulfills biosynthetic tasks in the production of branched-chain and surface exposed lipids, which are essential virulence factors in *Mtb* [16].

If the predominant MCIT pathway is lost, for example by genetic disruption, the MMCO drain in to lipid biosynthesis can re-route flux from a damaged MCIT pathway and relieve cytotoxic stress from MCIT intermediates [16]. Similarly, the MMCO pathway can partly compensate for the loss of MCIT flux up to a certain capacity, but only if its activity is induced [10, 36]. These findings indicate some degree of interplay and compensation between the two pathways, but quantitative and mechanistic understanding thereof have been lacking.

In this study we address the question why mycobacteria keep two seemingly redundant pathways for the same task of propionate catabolism, by means of thermodynamic-kinetic modeling, integrating absolutely quantified proteomics and metabolomics data. We explain why both pathways are required and why the bacteria favor the MCIT pathway despite the danger of self-intoxication by its intermediates.

The flux through a reaction or pathway is shaped by the expression of enzymes but also by metabolite dependent enzyme saturation and reaction thermodynamics (Figure 1 B, [7, 8, 11, 17, 24]). Sub-optimality in any of those factors in the cellular context can render a pathway unfavorable. Here, we use modeling approaches to test the significance of each of the factors for the two pathways in question. We find that each pathway is optimal in a different nutritional condition, based on pathway thermodynamics and the optimal distribution of protein masses. While these insights are revealed under steady state conditions, the full mechanistic interplay between the pathways requires to venture in to the dynamic time scale. With the help of a full thermodynamic-kinetic model based on fast nutrient switch experiments, we characterized the function of both pathways in detail: While the MMCO pathway can quickly react to sudden changes in the propionate concentration and detoxify the cell accordingly, the MCIT pathway acts as a professional catabolizer, which has a higher overall flux capacity.

## Results

### Thermodynamic driving forces are comparable between the two pathways in catabolic direction but prohibit biosynthetic flux through the MCIT pathway

Thermodynamics are a major flux shaping factor within cellular reaction networks [9, 11, 19, 24, and others]. The direction and flux efficacy of a reaction is restricted by its thermodynamic gradient, which is characterized by the Gibbs Energy of the reaction ∆*_r_G′*. Only if ∆*_r_G′* is negative, the reaction can proceed (*i.e.* has a positive net-flux). A lower ∆*_r_G′* thereby indicates a more favorable reaction, *i.e.* less effort is required to achieve a higher net-flux through the reaction. The value of ∆*_r_G′* depends on the reaction specific Standard Gibbs Energy ∆*_r_G*^0^ as well as on the ratio of substrate to product concentrations. Within a pathway, the ∆*_r_G′*s of all reactions have to be feasible, to allow flux. The reaction with the highest ∆*_r_G′* (*i.e.* the lowest absolute) is limiting the pathway flux, it acts as a thermodynamic bottleneck. One method to identify and quantify such bottlenecks is the calculation of the maximum-minimum driving force (MDF, [24]). It identifies the pathway reaction with the highest ∆*_r_G′*, given sensitive, physiological bounds on the metabolite concentrations within the pathway (for details see Supplementary Text 1.2).

To test whether *in vivo* the MCIT pathway might be preferred over the MMCO pathway due to less stringent thermodynamic bottlenecks, we first analyzed the thermodynamic gradients within the two pathways. We acquired absolute metabolite measurements in the model organism *M. bovis* BCG in conditions where either propionyl CoA catabolism (growth on propionate containing media) or biosynthetic flux towards propionyl CoA (growth on glucose containing media) was required. For both pathways we calculated the optimum driving force profiles and identified the limiting reaction with the highest ∆*_r_G′* (Figure 2 A).

**Figure 2:**
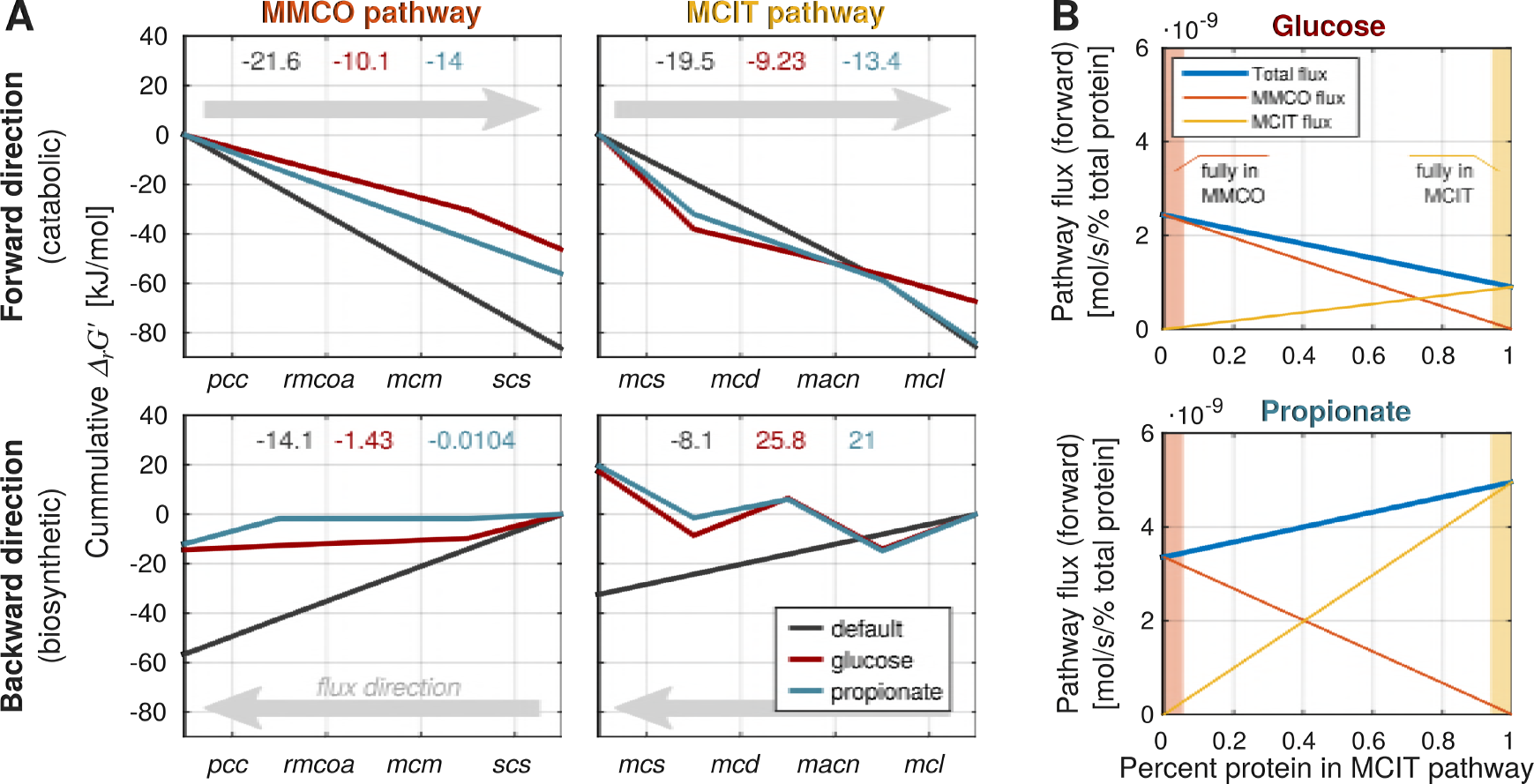
Optimum thermodynamics and protein investment. **A** Driving force profiles after maximum-minimum driving force optimization. The cumulative Gibbs energy ∆*_r_G′* along the pathway is plotted, the MMCO (left) and the MCIT (right) pathways are analyzed in forward/catabolic (upper panels) and backward/biosynthetic (lower panels) direction as indicated by light gray arrows. The participating reactions are labeled on the x-axis. Monotonously decreasing lines in the direction of flux imply pathway feasibility, increasing lines mark infeasible reactions. The different sets of bounds on the metabolite concentrations are shown in different colors: gray - default physiological bounds, red - measured concentrations under glucose conditions, blue - measured concentration under propionate conditions. Color coded numbers in the top of the plots indicate the ∆*_r_G′* of the bottleneck reaction in kJ/mol, positive values indicate pathway infeasibility. **B** Maximum pathway flux for limited enzyme availability. We allowed 1% of the total *M. bovis* BCG protein mass to be distributed between the two pathways in different ratios (x-axis, relative to the MCIT pathway). Hence, the bounds of each plot represent the full amount of protein being contained in one of the pathways (highlighted in the respective colors). The plots show the contribution of the MMCO (orange line) and the MCIT (yellow line) to the total flux (blue line) for each ratio (11 sampling points, *c.f*. Supplementary Figure S.1) for the growth on glucose (top) and propionate (bottom).

We found that while the thermodynamic profiles of the pathways differ, they have comparable bottleneck ∆*_r_G′* values in the catabolic direction (Figure 2 A, upper panel). In contrast, only the MMCO pathway is able to carry flux in the biosynthetic direction in both conditions (Figure 2 A, lower panels). The most strongly prohibiting reaction of the reverse MCIT pathway is the final splitting of methylcitrate into PCO and oxaloacetate by the *mcs* reaction.

### Optimum enzyme investment between the pathways is condition dependent

The amount of enzyme required in a pathway to achieve a certain flux is crucial for the efficiency of metabolic processes within a cell and its minimization is often considered an evolutionary objective. From an enzyme investment point of view, some bottlenecks might therefore not be as prohibiting as others. If the bottleneck reaction is catalyzed by a small protein which is easily synthesized, the cell can increase the flux through that pathway by producing additional enzymes at low cost. To account for this dependency on enzyme masses, we calculated the maximum flux through both pathways, given a limited total protein mass that can be distributed, as an evolutionary constraint, between the reactions of the two pathways. We focused on the catabolic direction, in which both pathways can carry flux.

We optimized the total catabolic flux through both pathways in the two metabolic backgrounds by scanning a range of possible ratios of enzyme mass distribution between the pathways (Figure 2 B). For each of the ratios, the total protein mass per pathway was fixed but could be distributed optimally between the pathways enzymes to achieve maximum flux (for implementation details see Supplementary Text S.1.3).

The analysis reveals that pathway optimality, considering thermodynamics as well as enzyme masses, depends on the metabolic background: In glucose media, the MMCO pathway can achieve a higher flux per enzyme mass (Figure 2 B, upper panel), whereas the MCIT pathway is preferable in propionate media (Figure 2 B, lower panel). Each of the pathways is therefore beneficial for the bacteria depending on the available nutrient. If the supply in nutrients is variable in the bacterial environment, it can be beneficial to retain both pathways’ in the genome, despite the higher biosynthetic costs.

### Dynamic detoxification of propionate pulses is predominantly facilitated by the MMCO pathway

Whereas the analyses of thermodynamics and optimal enzyme investment strategies revealed first insights into the characteristics of the two pathways, they largely neglect kinetic aspects of enzyme action. Firstly, they assume - due to the lack of available kinetic data - uniform catalytic parameters for each pathway enzyme. Secondly, both approaches are limited to the analysis of steady state behavior. However, in the changing environment of a ructious host cell, dynamic time scales are likely to be of importance in the understanding of metabolic functionality.

To characterize the dynamic behavior of the two pathways, we perturbed *M. bovis* BCG cells experimentally by a fast shift between the two metabolic conditions: From steady state growth on propionate to glucose media, in which reverse flux through the pathway in the biosynthetic direction will be required, and from growth on glucose to propionate media, where a fast detoxification of rising propionyl CoA levels is needed. We measured the dynamic metabolite concentration changes of both pathways at several time-points within the first 160 seconds after the switch (Figure 3, dots). Furthermore, we absolutely quantified the protein concentrations of the involved pathway enzymes in both culture conditions (propionate and glucose), which can be safely assumed to be constant over the short period of the shift experiment. We then built a full dynamic model (for details see Methods and Supplementary Text S.2) with thermodynamically consistent rate laws that include the major flux shaping factors (see Figure 1 B) to explain the observed behavior in detail. The model was calibrated to the dynamic metabolomics data and included also the measured protein concentrations for each nutritional condition. It was able to reproduce the experimental behavior (Figure 3, lines and shaded areas as error estimates, corresponding fluxes are shown in Supplementary Figure S.3)

**Figure 3:**
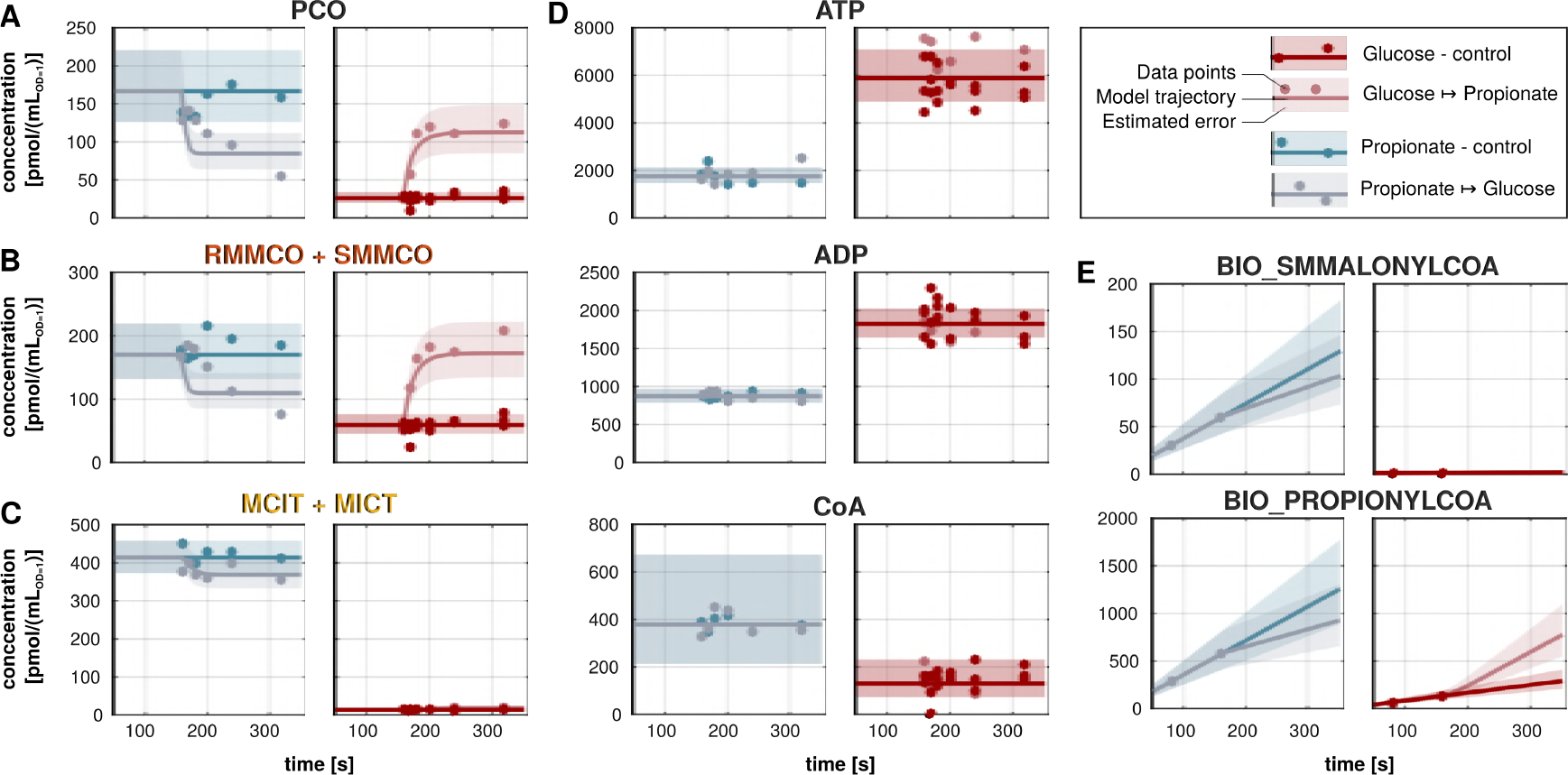
The MMCO pathway reacts dynamically to changes in the propionate supply, while the MCIT pathway is up-regulated in catabolic conditions. Experimental metabolomics measurements (dots) along with dynamic model simulations (lines) and the estimated error (shaded areas) for different nutrient switches at time t=160s. The model was fitted to the data shown here (values in Supplementary Table S.5) and used without further calibration for the predictions in all following simulations. **A** Propionyl CoA (PCO) is the initial metabolite of the MMCO and MCIT pathways and represents the entry point of propionate into the central carbon metabolism. **B** Observable of intermediate species of the MMCO pathway **C** Observable of intermediate species of the MCIT pathway (MCIT + MICT). **D** Co-factors, which are held constant in the model. **E** Dummy biomass species required to guarantee appropriate fluxes draining into biomass production. Please note that the final steady states after 160s depend on the enzyme concentrations (see Figure 1B), which are different in glucose and propionate conditions (Supplementary Table S.3).

Indeed, the two pathways show very distinct dynamic reactions to the nutrient shifts. Only the MMCO pathway (Figure 3 B) reacts dynamically to sudden increases and decreases in the propionyl CoA concentration (Figure 3 A). The MCIT pathway shows only a minor dynamic reaction, but its overall metabolite levels strongly change with the carbon source (Figure 3 C). We therefore conclude that the MMCO pathway facilitates the dynamic detoxification of propionate pulses.

### Thermodynamics limit MMCO flux

Our calibrated model further allowed us to analyze functional flux distributions in detail in order to identify the limiting factors for each pathway and condition. Using the model, we simulated metabolic challenges as they are likely to occur in the host phagosome, such as sudden changes in the availability of nutrients. These simulations enabled us to understand the specific metabolic tasks and limitations of the two parallel pathways.

We tested how changes in propionyl CoA concentration would impact on pathway fluxes in a given metabolic background. To this aim, the rate dependencies on the substrate concentration (similar to classical Michaelis-Menten curves) of the first enzyme of each of the two pathways were examined (Figure 4 A), already indicating the benefits and limitations of each pathway. The modular character of the thermodynamic rate law also allowed us to dissect the contributions of changes in enzyme concentration, enzyme saturation and thermodynamic limitations to the curve (Figure 4 B). To avoid a bias due to a local optimum of the parameter estimation, we calculated the mean curve of all feasible (**LL** < **LL***_min_* + ∆_95%_) parameter sets found during the calculation of parameter profile likelihoods ([30], Supplementary Figure S.4).

**Figure 4:**
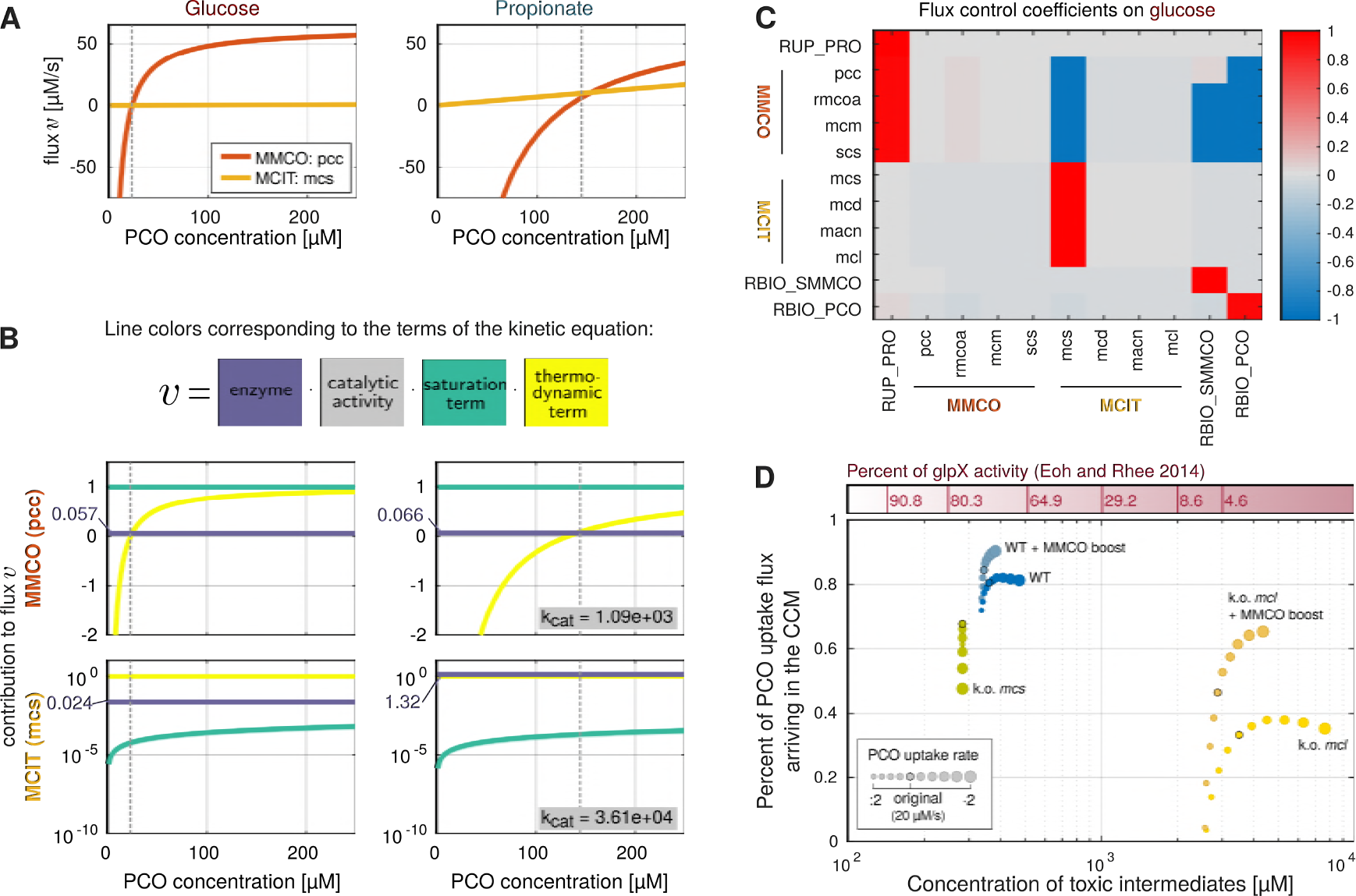
The trade-off between flux capacity and intermediate toxicity. **A/B** Substrate dependency of the first reaction step of both pathways dissected. The two experimental backgrounds glucose (left panel) and propionate (right panel) are shown with the vertical dashed line highlighting the mean PCO concentration of the metabolomics measurements. All simulated values are means of all feasible parameter sets of the profile likelihood estimation. **A** Michaelis-Menten-like curves of pcc (orange) and mcs (yellow), the first reaction steps of the MMCO and MCIT pathways, respectively. The dependency of the flux on the concentration of PCO is plotted. Note that the flux can become negative owing to the reversible nature of the thermodynamic rate law. **B** The total flux as depicted in A, decomposed in the contributions of the individual terms of the reaction rate (see also Figure 1 B): Enzyme concentration as measured by proteomics (dark blue, for clarity values in *µ*M are shown on the axes), kinetic term representing the saturation state of the enzyme (green) and the thermodynamic term quantifying the extent of the backward flux (yellow). The thermodynamic and the kinetic term are bounded by 1. The product of the three terms and *k^cat^* (lower right corner, right panel, given in 1/s) will give the values plotted in A. Upper panel shows dissection for the MMCO reaction pcc, the lower for mcs from the MCIT pathway. Note that the MCIT terms are plotted on a logarithmic scale. **C** Mean flux control coefficients in glucose media. The color coded (red - positive control, blue - negative control, intensity of the color encodes strength), normalized flux control coefficients are shown. The reactions labeled on the x-axis thereby exert the control on the fluxes through reactions shown on the y-axis. **D** Simulation of mutant strains. Varying propionate input fluxes (2 fold increase and decrease) were simulated for five different strains in a propionate media background. The trade-off between the accumulation of toxic intermediates and the percentage of carbon that is channeled into the central metabolism ("flux capacity”) is shown. For comparison, the toxic effect of the intermediates on fructose 1,6-bisphosphatase (in % remaining activity of the enzyme) as measured by Eoh and Rhee [10] is depicted in the top bar. Please note that those simulations were only carried out with the optimum parameter set.

From the rate dependency curves it becomes evident that under biosynthetic conditions (glucose consumption, Figure 4 A and B, left), the MMCO flux (orange) tightly follows the PCO concentration, with the potential to quickly switch from negative (biosynthetic) to positive (catabolic) rates around the measured PCO concentration (gray dotted line). This switch is solely facilitated by the adaptation of the thermodynamic gradient of the reaction (Figure 4 B, upper left, yellow line). The flux through the MCIT pathway is almost insensitive to changes in the PCO concentration as the first enzyme is strongly under-saturated at the lower PCO levels (Figure 4 B, lower left, green line). Hence, a sudden PCO pulse, as in our experiments, will be solely drained via the MMCO pathway, which rapidly shifts its reaction direction.

Under catabolic conditions (propionate consumption, Figure 4 A and B, right), the MCIT reaction has a higher capacity due to a drastically increased enzyme level (Figure 4 B, lower right, dark blue line) and hence out-competes the MMCO reaction. In the catabolic background, the *pcc* reaction has a shallow thermodynamic gradient (Figure 4 B, upper right, yellow line), most likely due to the decreased level of the co-substrate ATP. It would even invert its direction upon a slight decrease in the PCO concentration, consequently not fulfilling any catabolic tasks. Here, as well as in the biosynthetic case, the MMCO flux quickly saturates at increasing PCO concentrations in the physiological range, explaining the upper capacity bound for the pathway observed experimentally [10, 36]. The corresponding MCIT enzyme is still in the linear regimen of the rate dependency curve, guaranteeing a proportional change of the flux with the PCO concentration.

### Metabolic control is executed mainly by the first enzyme of the MCIT pathway

Employing metabolic control analysis [14] we could also resolve why the bacteria tune the metabolic flux distribution by up-regulating *prpC*, the enzyme of the methylcitrate synthase reaction *mcs* in the MCIT pathway. Metabolic control analysis is a standard method to quantify the influence of model entities, such as enzyme levels or individual parameters, on the system’s steady state. As we see from the flux control coefficients, which describe the effect of a local change in one enzyme concentration on the steady state pathway flux, the only significant control under biosynthetic conditions is exerted by *mcs* (Figure 4 C). Hence, only an up-regulation of *mcs* can significantly increase the steady state catabolic flux as required when propionate is to be used as a carbon source. The flux through MMCO reactions is only passively following the propionate uptake rate (RUP_PRO). Under catabolic conditions the control shifts also to the reactions of the MMCO pathway (Supplementary Figure S.6), consistent with the pathway’s compensating function described.

### Quantitative assessment of the trade-off between flux capacity and intermediate toxicity

We conclude that the thermodynamic properties allow the MMCO pathway to react quickly to changes in the PCO concentration, but also limit its overall capacity, rendering it insufficient to supply enough carbon for bacterial growth. The MCIT pathway, on the other hand, gains momentum only if its enzyme expression is increased, thereby reaching a higher capacity and avoiding saturation effects to efficiently channel carbon to the central carbon metabolism. As a drawback, this pathway contributes to the risk of accumulating toxic intermediates.

We used the kinetic model to quantitatively asses how this trade-off between toxicity and high flux capacity is efficiently balanced by the two pathways. We thus simulated increased and decreased carbon uptake via PCO compared to the reference condition in the model to quantify which percentage of the carbon is fed into the central metabolism (“flux capacity”) where it can be used to generate biomass precursors and energy. At the same time, we monitored the absolute concentrations of toxic intermediates. With the help of *in-silico* knock-outs we further dissected the interplay between the two pathways in this trade-off (Figure 4 D). For comparison, phenotypes of related experimental knock-out studies are summarized in Supplementary file S3.

In the wild type (dark blue) increasing influx is only transmitted via both pathways up to a certain capacity (~80%), the rest is drained into lipids. Further increase in the propionate uptake results in raising levels of toxic intermediates, but to levels that are only mildly inhibiting fructose 1,6-bisphosphatase [10]. A boost in the MMCO pathway activity (light blue, here done by increasing the activity of *rmco* 100-fold, the reaction with highest control coefficient in the pathway) can both lower toxic metabolite levels and increase flux capacity to ~90%. This additive behavior of the two pathways is consistent with experimental observations [36]. The capacity limitations in the MMCO pathway become evident in *in-silico* mutants with a disrupted MCIT pathway. In the also experimentally well studied *mcl* knock-out, the MMCO pathway can only achieve ~40% flux capacity in our simulations (bright yellow). If boosted as described above, it can reach up to ~65% capacity (orange) - still much less than the wild type. Importantly, the MMCO boost cannot rescue the intermediate toxicity in our simulations. If the first enzyme of the MCIT pathway *mcs* is lost (green) the compensation by the MMCO pathway is even lower, with flux capacity decreasing below 50% with increasing input flux, accompanied by a drastically increased buffering flux into lipids. However, these cells do not experience intermediate toxicity, in accordance with experimental studies [45, 43].

Taken together, our simulations explain the metabolic capabilities of the two pathways in a quantitative manner. The trade-off between flux capacity and intermediate toxicity is thereby identified as a structural weakness of the catabolizing system that can be exploited to fight *Mtb* infection.

## Discussion

### Professional high-capacity catabolism vs. flexibility for rapid detoxification

The presented analyses characterize the MCIT pathway as a high-capacity catabolic pathway that exploits thermodynamically beneficial reaction steps, which operate below saturation in physiological scenarios. The MMCO pathway acts as a flexible pathway capable of rapid PCO detoxification but also of biosynthetic tasks. This flexibility is facilitated by a fine-tuned thermodynamic balance at the cost of an upper catabolic flux capacity limit.

These distinct functionalities render it beneficial for the cell to retain both pathways despite their redundancy in channeling carbon from PCO to the central carbon metabolism. Implementing both in one pathway would reduce the efficiency of either functionality, as a high forward flux capacity with a steep thermodynamic gradient contradicts the ability to quickly invert the flux direction.

Further metabolic tasks of each pathway could be seen as circumstantial of the described functionalities. The constitutively expressed MMCO pathway also supplies precursors for virulence relevant lipids, which are required under all environmental conditions. The MCIT pathway, as an additional metabolic advantage, directly feeds pyruvate into lower glycolysis, thereby skipping potentially expensive gluconeogenetic reactions.

### Implications for treatment strategies

The MCIT pathway has been subject of extensive study as an anti-tubercular drug target, encouraged by the lack of a homologue pathway in the human host. Recently, also its role in antibiotic tolerance by specifically rerouting central carbon metabolic fluxes was described [23]. The pathway also occurs in other pathogens and is up-regulated upon exposure to propionate, which is often part of their diet in the pathological context *(Salmonella enterica* [28], *Toxoplasma gondii* [18]). The plasticity of this crucial lipid biosynthetic and catabolic node has recently been shown to be relevant in eukaryotes, with implications on human metabolic diseases [46], arguing for a general metabolic strategy. Functional understanding of the pathway functionality and its interplay with the compensating MMCO pathway as presented in this study is highly relevant for the development of new treatment strategies in these contexts.

Eoh and Rhee [10] as well as Savvi and coworkers [36] show that in mycobacteria the MMCO pathway can compensate for the loss of MCIT pathway functionality in ∆*icl* mutants in terms of survival but cannot support growth in an equally efficient manner. However, Van der Ven *et al.* [45] and Upton and coworkers [43] pointed out that the inhibition of the upstream enzyme *prpC* can reverse this effect, implying that the growth defect is due to the accumulation of toxic MCIT intermediates rather than due to a less efficient metabolic processing of carbon through the MMCO pathway. Our mutant simulations quantitatively underline these findings.

Consequently, the optimum bactericidal strategy would be to inhibit both a downstream MCIT enzyme such as *icl* as well as the first enzyme of the MMCO pathway, thereby increasing toxic intermediate levels and abrogating the buffering flux bypass completely.

### Quality of model calibration

Despite the limitations of modeling approaches - here for example the inability to resolve the dynamics of propionate decay with high confidence - mechanistic conclusions can be drawn from the presented analyses. Models can only yield predictions up to the level of accuracy and coverage of the data. But during the modeling process, we can pinpoint parts of the system which are still ill-defined, for example by identifying parameter non-identifiabilites [30] which were present for the majority of the model parameters, despite the high number and quality of data points included. However, other parts of the model are well calibrated (identifiable parameters). Furthermore, mechanisms can become evident for all possible scenarios of the non-identifiable parts of the model. For this reason, we performed our simulations with all sets of parameters identified as yielding a model quality below the likelihood threshold in the profile likelihood estimation. Our simulations therefore cover a range of feasible behaviors given the experimental data, not just an individual point in the parameter space.

Generally, dynamic metabolic modeling efforts often face a lack of measured kinetic parameters, especially in the appropriate condition. *in vitro* experiments investigating enzyme kinetics are sparse for most pathways and organisms, always assay dependent and never reflect the intracellular situation. In contrast, model calibration to absolute intracellular metabolite and protein concentrations can determine enzyme kinetics *in vivo* in the natural environment of corresponding metabolic pathways.

### Dynamics and quantitative data make the difference

The thermodynamic and enzyme cost analyses clearly highlight the condition dependent nature of pathway optimality: The cheapest pathway in one condition might be out-competed by an alternative route under different conditions. Despite their usefulness in many studies [8, 11, 17, 48], such approaches are limited by their focus on time scales that allow for equilibration of the metabolism for describing the evolutionary benefit of reducing resources and increasing metabolic outcome. However, as *Mtb* invest only a small fraction of their resources in proteins [3, 33], the evolutionary pressure of reducing protein cost might not be high for these bacteria. In addition, the cells might have other objectives for pathway functionality that are not detectable by optimizing for a constant condition, especially for bacteria subjected to variable environmental conditions as those *Mtb* faces in the human host. In this context, multi-objective Pareto optimality has been found to describe the overall behavior of systems well [8, 40]. Here, we add a distinct, temporal objective, describing metabolic non-steady state optimality. The transient behavior in response to sudden metabolic challenges has proven to be essential for the system in question here.

We also highlight the importance of concentration constraints based on quantitative metabolite measurements. For example, only the measured metabolite concentrations reveal the infeasibility of the MCIT pathway in biosynthetic direction, while under default physiological concentration bounds (1*µ*M-50mM) the MCIT pathway would falsely be classified as reversible (Figure 2 A, lower right, gray line). The actual intracellular concentrations are far more strict constraints on the feasible flux space, which underlines the importance of high quality metabolomics data to understand the underlying mechanisms.

In conclusion, our modeling-based approach was able to resolve the functional differences in the two seemingly redundant pathways responsible for catabolizing *in vivo* carbon sources. The combination of dynamic modeling and absolutely quantified experimental data revealed transient constraints in thermodynamics as well as intermediate toxicity that force the cell to retain both pathways, explaining an important aspect of metabolic plasticity in *Mtb*.

## Methods

### Experimental procedures

#### Cell culture and experimental setup

The experiments were conducted with *M. bovis* BCG, a widely used *Mtb* model system which requires lower biosafety level and shares 99.5% of the *Mtb* genome [1]. *M. bovis* BCGis less persistent and virulent than *Mtb* but many biological features were found to be similar (*e*.*g*. flux distributions [4], ability to grow on cholesterol [44]). In our experiments, *M. bovis* BCG was grown on Middlebrook 7H9 medium without glycerol containing exclusively propionate or glucose as carbon source (0.5 g/L). Samples were taken in exponential growth phase, transferred to a Büchner funnel with nitrocellulose filter. For the shift experiments, the cells were perfused with fresh media with the culture carbon source for 1 minute to allow acclimatization to the new aeration conditions. Afterwards perfusion media was changed to the respective other carbon source. The method is described in detail for mycobacteria in Murima *et al.* [22]. Additionally, proteomics samples were taken from the culture.

#### Proteomics measurements

For protein quantification, cells were disrupted and subjected to tryptic digestion. Resulting peptides were quantified by absolute label free quantification [38] or internal heavy isotope labeled protein standards [39], where available. Proteomics measurements are summarized in Supplementary Table S.3.

#### Metabolomics measurements

Cells on the filters were subjected to ethanol extraction after 0, 10, 20, 40, 80 and 160 seconds of perfusion with the new media and absolutely quantified by mass spectrometry by normalizing to ^13^C-labeled internal standards as described before [6]. Metabolomics measurements are summarized in Supplementary Table S.5. We selected the final time-point of 160s as the metabolite concentrations have reached a new steady state during that time, while we can assume negligible changes in the protein concentrations. This allows us to include them as constant in our model simulations.

### Modeling, simulation and optimization procedures

#### Compilation of pathway reactions

The reactions and the corresponding enzyme complexes of both pathways used for all modeling approaches were compiled from literature and are summarized in Supplementary Table S.2. The pathways were embedded in the larger biological context via boundary fluxes based on biomass backtracking [42] from a genome scale model of *M. tuberculosis* [33]. The compiled reaction were used for all simulations and optimization approaches as well as for the dynamic model (see below).

#### Thermodynamic constants

The Gibbs energy of a reaction is defined as 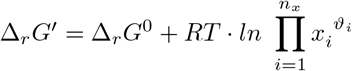. It can be calculated from the standard Gibbs energies of the reactions ∆*_r_G*^0^, the substrate and product concentrations *x_i_*, their respective stoichiometric coefficients *ϑ_i_*, the absolute temperature T and the ideal gas constant R. At ∆*_r_G′* = 0 the forward flux through an reaction equals the backward flux, such that the net-flux of the reaction vanishes. For our calculations, the standard Gibbs energies of the reactions ∆*_r_G*^0^ were obtained from the component contribution method (eQuilibrator tool [12, 26]) at pH 7.5 and a total ionic strength of 0.2 M, they are summarized in Supplementary Table S.3.

#### Nonlinear constraint based optimization

Optimizations were performed in Matlab using the fmincon optimizer that allows for the integration of linear and nonlinear constraints on the optimization problem. Optimizations were carried out with Latin Hypercube [20] distributed initial points and convergence was examined over a sufficiently large number of optimization runs. Mathematical formulations and detailed information on all optimization problems can be found in the Supplementary Text S.1. In all cases the default concentration bounds were between 1 – 50 *·* 10^3^*µM* (based on [2, 11, 50] and metabolomics data presented here).

#### Dynamic ODE modeling

The model reactions and their kinetics are summarized and explained further in Supplementary Table S.2. The enzymatically modeled reactions (shown as black arrows in Figure 1) were modeled using a decomposition of the Michaelis-Menten equation that differentiates between kinetic (saturation) term and a thermodynamic term (Equation in Figure 1 B, [25]). The uptake of propionate was modeled by a constant rate, perturbed by step functions representing the media switches. Cofactor (OXA, CoA, H_2_O, ATP, ATP, P*_i_*, CO_2_, PYR) concentrations were held constant, but their concentration value was estimated in accordance with the experimental values, where available. For non-measured cofactors the default concentration bounds were applied. Biomass flux was modeled as a simplified Michaleis-Menten kinetics linked to the overall biomass production by biomass backtracking [42] and corresponding dummy biomass observables (see Supplementary Text S.2). Enzyme concentrations in the kinetic laws were taken directly from the proteomics measurements of the two growth conditions and assumed to be constant for the comparably short time of the dynamic simulation after the nutrient switch (160s).

Overall the model has 94 parameters of which 34 are initial concentrations, 10 error model parameters and 8 Gibbs free energies (Supplementary Table S.1). The model comprises 14 reactions (Supplementary Table S.2), 9 dynamically modeled species and 11 constant inputs and co-factors (Supplementary Table S.4).

#### Computational implementation and parameter estimation

The model was implemented in the D2D modeling framework [31]. Experimental data was mapped via observables *y* (Supplementary Table S.4), a logarithmic error model of the form *y_m_ · σ_y_* was assumed and estimated alongside the model parameters.

The objective function consists of the negative log likelihood of the model *y_m_* given the data *y* and the error model and the fulfillment of applied steady state constraints. The constraints are necessary to guarantee the model to be in steady state before the media shift is simulated and were implemented as minimization of the metabolite trajectories’ derivatives at the initial point of the simulation (t = 0).

Optimizations were run with Latin Hypercube distributed initial points and the lsqnonlin optimizer in D2D.

#### Calculation of profile likelihoods

Profile likelihoods [30] were calculated to asses parameter confidence bounds and identifiability. They are summarized in Supplementary Figure S.4and Supplementary Table S.1.

#### Metabolic control analysis

The calculation of the metabolic control and response coefficients [14] was implemented within D2D, based on the symbolically calculated matrices for *∂v/∂x* and *∂v/∂p*.

#### Accessibility

Programming code (public git repository https://github.com/tbphu/pdtx upon acceptance, reviewer access: login - PdtxReviewerMSB, password - PdtxReviewerMSBPassword) as well as the models in SBML format (Supplement) are available online.

## Acknowledgements

We would like to thank Max Schelker for help with the D2D framework and Jannis Uhlendorf for help with implementing MCA in D2D. This project was supported by the European Commission project SysteMTb (HEALTH-F4-2010-241587).

